# Genomic repeatability and predictability of climate (mal)adaptation in a reef-building coral

**DOI:** 10.64898/2026.01.17.700041

**Authors:** Zoe Meziere, Andreas Bachler, Iva Popovic, Yves-Marie Bozec, Jennifer McWhorter, Christopher Coppin, Katrina McGuigan, Cynthia Riginos

**Affiliations:** Australian Institute of Marine Science, Townsville, 4810 QLD, Australia; School of the Environment, The University of Queensland, St Lucia, 4072 QLD, Australia; CSIRO Black Mountain Laboratories, Canberra, 2601 ACT, Australia; Rosenstiel School of Marine, Atmospheric, and Earth Science, University of Miami, Miami, FL, USA

**Keywords:** climate adaptation, genotype-environment associations, repeated evolution, coral genomics

## Abstract

Climate change is a growing threat to biodiversity, and the persistence of populations largely depends on their capacity to adapt to changing environmental conditions. Although there is an urgent need to forecast local adaptive potential, it is unclear how such predictions are affected by the genomic architectures underlying local adaptation across a species’ range. In this study, we examine the genomic basis of local adaptation of the short-distance dispersing coral *Stylophora pistillata*, sampled at forty-six sites across eight reefs of the Great Barrier Reef, Australia. Our results show that thermal adaptation for this species involves hundreds of genomic loci with combinations that differ across geographic regions. Although adaptive loci were largely region-specific, genotype-environment relationships estimated across the entire range could predict regional-level adaptive patterns. This shows that genome-wide sequence data combined with geographically broad sampling can support reliable evolutionary forecasting. Under climate change projections, predicted shifts in genotype-environment associations were highly spatially variable, both between and within geographic regions. While some populations might be sufficiently adapted for moderate (SSP1-2.6 and SSP2-4.5) climate warming by 2050, up to 30% may face severe maladaptation risk by 2100 under a high-emission (SSP5-8.5) scenario. Collectively, these findings offer new insights into the spatial distribution of coral adaptive potential and how it might shape corals’ resilience in a warming ocean.

## Introduction

Future climate projections put many species at risk of population decline, range contraction and extinction (1, 2), threatening entire ecosystems. While reducing greenhouse gas emissions is paramount for limiting biodiversity loss (3), fundamental science must also advance to provide evidence-based and evolution-informed conservation strategies (4, 5). Adaptive capacity is critical for long-term species persistence (6, 7), and as the climate warms, thermal adaptation will undoubtedly shape species’ evolutionary trajectories in the coming decades.

Because standing genetic variation can enable rapid adaptation (8, 9), the scope for adaptive change might be predictable from distributions of present-day genetic variation (10, 11). Local adaptation arises when divergent selection favours different genotypes in different environments (12, 13), with a species’ dispersal capacity (mean axial distance σ) and the strength of selection (*s)* setting the spatial scale at which it evolves, approximately 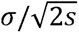 in a continuous two-dimensional habitat (14, 15). Species with limited dispersal can therefore maintain adaptive divergence over short geographic distances, even under modest environmental selection (16), with different genes possibly enabling convergent responses to similar environmental gradients across geographically distant populations.

Whether patterns of local adaptation are repeated — and thus predictable — across populations is a fundamental question in evolutionary biology. The same loci might be recurrently selected upon in populations facing comparable environmental contrasts when these populations also share similar genomic backgrounds or high standing genetic diversity (18). Conversely, selection may target different, population-specific loci when genomic backgrounds differ (8, 19, 20) or when beneficial mutations are highly redundant (21). Even without locus-level repeatability, the same pathways might still be involved in local adaptation (22). Genomic architectures may also inform how many genomic routes are available for adaptation (23, 24): adaptation from few large-effect loci (i.e., oligogenic) might be less redundant and more predictable, whereas adaptation via many small-effect loci (i.e., polygenic) might be more redundant and less predictable (18, 25). Understanding the genomic basis of local adaptation and its repeatability across populations is therefore essential for identifying adaptive loci and predicting a species’ adaptive potential.

Traditional approaches such as reciprocal transplantation and common garden experiments (13), offer great insights into local adaptation, but are impractical for many species and require prior knowledge of fitness-related traits (26). Genotype-environment associations (GEAs) offer an alternative approach, using correlations between genomic and environmental variation to identify candidate loci and infer adaptive clines across a species’ range (27–29). However, these tests are prone to false positives when environmental gradients align with population structure (30–32) and are often performed across large geographic extents (e.g., spanning hundreds to thousands of kilometres) that may mask adaptation dynamics occurring at finer spatial scales (33).

Contemporary GEA patterns can also be used to predict maladaptation under future environmental conditions (i.e., genomic offset, reviewed in 29). In practice, if local adaptation is predictable in space and time, such models could inform management by identifying pre-adapted populations with genetic attributes that match projected conditions (34). However, genomic offset models implicitly assume GEAs represent range-wide patterns, and if the repeatability of GEAs across a species’ range is low, genomic offset predictions may be unreliable. This assumption has rarely been tested, and is needed to assess the robustness of GEA and genomic offset methods under realistic empirical situations (35).

Most genomic studies of climate adaptation and its predictability have focused on terrestrial plants (36), while marine species have received comparatively little attention. Corals are highly susceptible to rising ocean temperatures (37, 38), and as foundational species, their survival underpins the functioning of entire reef ecosystems. Coral species are likely under strong selection for heat tolerance and studies have demonstrated a heritable basis for temperature-related traits (e.g., 39, 40), with dozens to hundreds of associated candidate loci identified via genome scans (reviewed in 41). Although we expect polygenic architectures to be more redundant (25), particularly for dispersal-limited corals, this has not yet been evaluated. Thus, it remains unknown whether — and to what extent — the genomic basis of thermal adaptation varies across coral species’ ranges.

Coral reefs are highly heterogeneous environments, making coral species tractable systems for studying the repeatability of local adaptation among reefs. Steep gradients in temperature, light, water flow, and turbidity occur over meters to tens of meters, driven by complex bathymetry and hydrodynamics (e.g., 42, 43). For example, reef flats and lagoons experience more variable temperatures, and wave-sheltered habitats are typically warmer than wave-exposed habitats (44, 45). Along a reef slope, temperature and light availability also change steeply with depth — often more steeply than they do across equivalent elevation changes on land. For short-dispersal corals, this fine-scale environmental variation is expected to favour micro-geographic local adaptation within reefs (16). Unlike plants and trees, where decades of provenance trials and transplant experiments have generated rich datasets on local adaptation (46), such multigenerational experimental resources do not exist for corals. Nonetheless, standardised heat stress essays (e.g., 47–49), longer-term common garden experiments, (e.g., 50), and reciprocal transplants between reef patches (e.g., 51, 52) or depths (e.g., 53) have documented fitness trade-offs, home-site advantage, and persistent physiological differences among conspecific corals inhabiting adjacent reef habitats, consistent with within-reef adaptation.

The brooding coral *Stylophora pistillata* is an ideal system for testing parallel adaptive responses. With meter-scale larval recruitment (54), dispersal does not exceed the extent of environmental heterogeneity among reef habitats, and local adaptation may have evolved at fine spatial scales in this species (16). As a result of limited gene flow among distant reefs, strong genetic differentiation has evolved among regional populations on the Great Barrier Reef (54). Here, we combined environmental and whole genome data of *S. pistillata* to (i) assess how repeatable the genomic basis of thermal adaptation is across reef populations, (ii) evaluate the predictability of adaptive genomic patterns to a multivariate environment, and (iii) investigate how these results affect genomic forecasting under climate change.

## Materials and Methods

### Whole genome re-sequencing

237 *S. pistillata* coral colonies were sampled from eight reefs across the Great Barrier Reef (GBR): Lady Musgrave Reef, Heron Reef, the Central Reefs (Little Broadhurst, Chicken and Davies), Pelorus Reef, Moore Reef and Lizard Reef (Figure 1A, Figure S1). These reefs are geographically distant, spanning more than 11 degrees of latitude. At each reef, we sampled between 2 and 6 sites (Table S1), with adjacent shallow (4-7 meters) and deeper sites (12-15 meters) at most locations (Figure 1A, Figure S2). We sampled between 8 and 114 coral colonies per reef (Figure S2, Table S1). This hierarchical sampling design allowed us to capture environmental variation between reefs but also within reefs (Figure 1C), with a total of 46 sites and between 1 and 12 samples collected per site (Table S1). These samples were initially collected as part of a larger dataset analysed using reduced-representation sequencing (55). In this previous study, we discovered that *S. pistillata* is a species complex, comprising five genetically distinct and sympatric species in the GBR. In the present study, we re-sequenced all samples assigned to *S. pistillata* Taxon1 using whole-genome sequencing. Details related to field collection and sample preservation can be found in (55).

**Figure 1.**
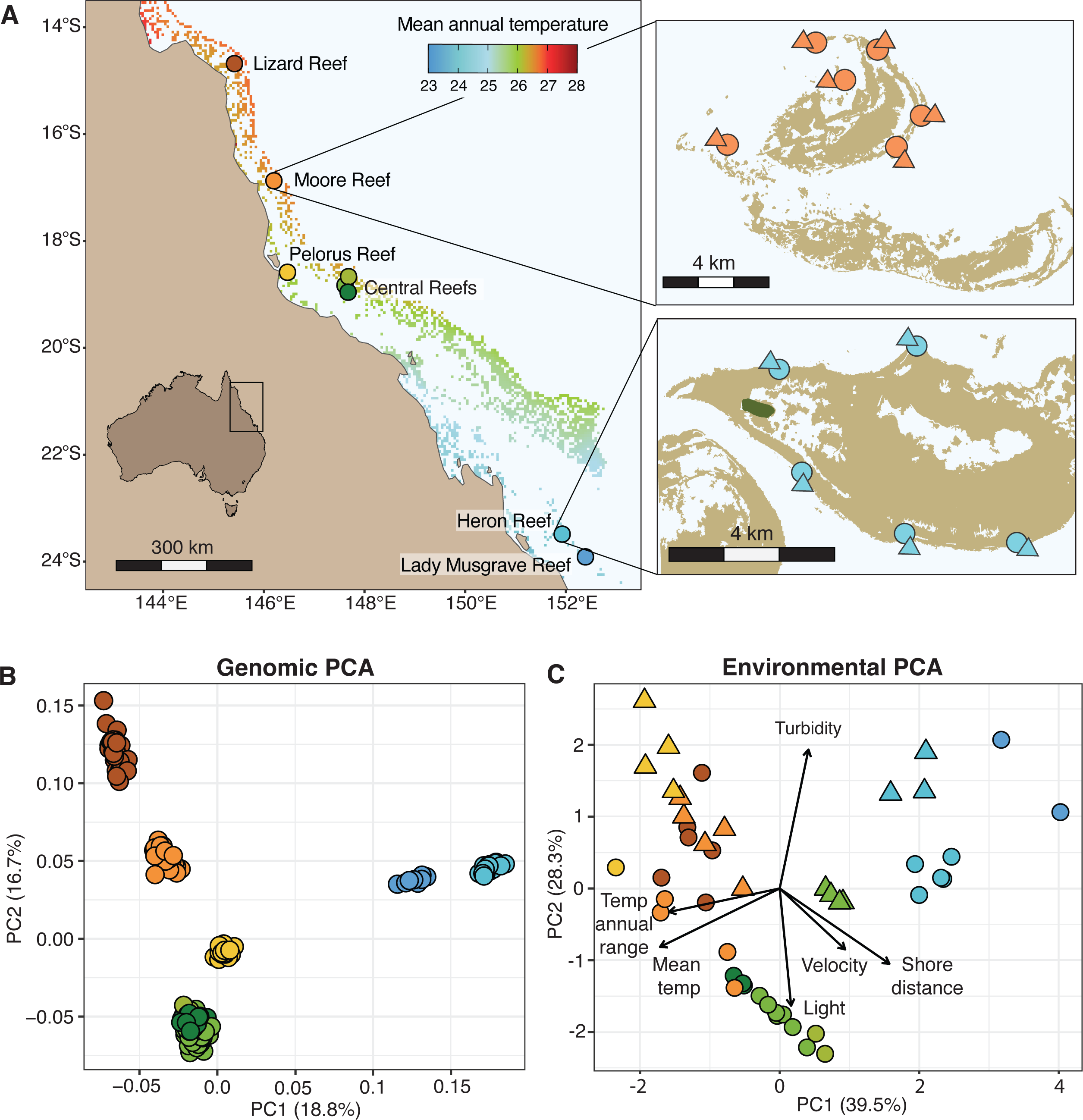
Sampling design, population structure and within-reef environmental variation. (A) *Stylophora pistillata* coral colonies were sampled at eight reefs located along a latitudinal temperature gradient on the Great Barrier Reef, Australia. Temperature gradient represents mean annual temperature from NOAA Coral Reef Watch between 1985 and 2024. Davies, Little Broadhurst and Chicken Reefs are grouped as ‘Central Reefs’. Insets show two examples of the nested sampling design where multiple sites were sampled at each reef, including adjacent shallow (4-7 meters, circles) and deeper sites (12-15 meters, triangles); (B) Principal Component Analysis reveals strong reef-level population genetic structure based on 1.3 million SNPs. Samples are shown in the same colours as their corresponding reef locations in panel A; (C) Principal Component Analysis summarising environmental gradients across sampling locations. Sites are depicted with the same colours and symbols as in panel A. Environmental variables include temperature mean, temperature annual range, light intensity, turbidity, wave orbital velocity at the seabed and distance to shore (eReefs, 2015-2023).

Genomic DNA was extracted using the QIAGEN Blood and Tissue kit following manufacturers recommendations. Whole-genome libraries were prepared using the Lotus DNA Library Prep Kit for NGS with 10 ng of input DNA and enzymatic fragmentation to achieve average insert sizes of 350 bp. Final amplification consisted of 8 PCR cycles and libraries were quantified using a Qant-iT dsDNA assay kit and libraries were multiplexed in equimolar ratios prior for sequencing. Sequencing was performed by Azenta Life Sciences (Suzhou, China) on a NovaSeq 6000 S4 flowcell (300 cycles, 150 bp paired-end reads).

### Read processing and variant calling

We used FASTQC (http://www.bioinformatics.bbsrc.ac.uk/projects/fastqc) and MultiQC (56) to examine raw read quality and adapter contamination. Basic filtering of raw reads was performed in Trimmomatic (57) by keeping bases with a minimum phred-score quality of 20 with a sliding window of 4 bp, removing adapter sequences with standard Illuminaclip parameters and removing reads shorter than 50 bp after trimming. Trimmed paired reads were then mapped to the *S. pistillata* reference genome (GenBank assembly GCA_032172095.1) using the Burrow-Wheeler Aligner (58) and the BWA-MEM algorithm with default settings. Resulting SAM files were converted to sorted and indexed BAM files using Samtools v1.10 (59). We assigned read groups to BAM files and removed PCR duplicates using picard (http://broadinstitute.github.io/picard/).

We followed a standard GATK pipeline to call variants (60). Specifically, we used GATK HaplotypeCaller for each individual and consolidated resulting GVCF files into a Genomics Database using GATK GenomicsDBImport. We then performed joint genotype calling across all samples for each contig using GATK GenotypeGVCFs and gathered contig-specific variant files into a single VCF file using GATK GatherVcfs. We selected Single Nucleotide Polymorphisms (SNPs) only using SelectVariants.

The dataset (‘all individuals’) was further filtered using VCFtools v0.1.16 (61) with the following parameters: *--min-alleles 2, --max-alleles 2, --remove-indels, --mac 3, --minQ 30, --minGQ 20, --min-meanDP 10, --max-meanDP 20, --max-missing 0.8*. Additionally, the ‘all individuals’ dataset was split into population datasets to perform population-level analyses. Populations were chosen based on PCA groupings assigned below. Each population dataset was then filtered using the same parameters as above.

To investigate population structure across the sampling range, we first used PLINK v2.0 (62) to remove SNPs in linkage disequilibrium (LD) from the ‘all individuals’ dataset (*--indep-pairwise 200 20 0.5*) and then performed a Principal Component Analysis (PCA).

### Acquisition of environmental data for GEA analyses

To obtain environmental data for each sampling site, we first extracted environmental variables from the eReefs GBR1 and GBR4 hydrodynamic models (63) with a monthly time step, specifying the date range 01/2015 - 12/2023 (108 months total), 5 meters depth for the shallow sampling sites and 13 meters depth for the deep sampling sites. We also extracted 10m-resolution wave exposure data using estimates generated by (64) who computed wave data from long time series wind data using the Simulating Waves Nearshore (SWAN) model. We investigated multi-collinearity among all variables using Pearson’s correlation coefficients (Figure S12). We retained six variables that share less than 50% of their variation (Pearson’s coefficient r<0.7, equivalent to r²<0.49, Table S4) and that have previously been shown to predict coral geographical distributions (65–67): mean temperature, temperature annual range (calculated as the difference between maximum monthly mean and minimum monthly mean), mean light intensity (calculated as photosynthetically available radiation, PAR), mean turbidity, mean orbital velocity on the seabed (a measure of wave energy) and distance to shore. These variables were used differently in subsequent analyses. We used mean temperature in a univariate framework to investigate repeatability of adaptive signatures among populations, as there is evidence for fine-scale thermal adaptation in other coral species. We then used all six variables in a multivariate framework to investigate the predictability of genome-wide adaptive signals across populations.

### Genomic signatures of temperature adaptation within populations

To investigate signatures of local thermal adaptation, we estimated correlations between allele frequencies and mean temperature for each population (i.e., genomic PCA cluster) independently. One population (Lady Musgrave Reef) was excluded from population-level analyses due to small sample size (see Results). First, we converted individual-based data into allele frequency data by combining individuals from the same sampling site and calculating per sampling site allele frequencies using VCFTools. Although this increases the risk of allele frequency sampling error when only few individuals are sampled at a given site, it increases power in downstream GEA analyses by reducing individual-level variability. Then, for each population-level dataset, we correlated site-level allele frequencies with historical annual mean temperature using Kendall’s τ (Supplementary Information). Temperature data derived from hydrodynamic models is at a coarse resolution despite being the best data available for this study. It is therefore possible that fine-scale thermal differences among sampling sites were not captured in this analysis, which would result in an underestimation of the number of adaptive loci. Using temperature loggers at each location was outside the scope of this study.

We then used the Weighted-Z Analysis (WZA) approach (68) (Supplementary Information) to combine correlation information in 31,828 10-kb unique 10-kb genomic windows. This window size was based on LD decay and gene length (Supplementary Information, Figure S13, Figure S14) but the analysis was also conducted using 20-kb windows for comparison. WZA was run for each population using Kendall’s τ *p*-values, removing SNPs with minor allele frequency below 0.05. Kendall’s τ *p*-values were converted into empirical *p-*values, which were used to calculate window-level weighted-Z scores (68). To account for variation in SNP density per genomic window, we down-sampled the most SNP-rich genomic windows to the 75th percentile (mean of 77 SNPs/window across datasets) and calculated the average weighted-Z score across 100 iterations (68). We also required a minimum of three SNPs per genomic window, following the standard parameters described in (68). To evaluate whether the observed genotype-environment associations were stronger than expected by chance, we performed permutation tests (Supplementary Information).

We identified candidate adaptive genomic regions exceeding the 99^th^ percentile of WZA empirical *p*-value (i.e., top 1% most significant windows) within each population. Ranking windows was preferred over an arbitrary *p*-value threshold, because empirical *p*<0.05 did not distinguish outlier windows from background signal (Figure 2). This is aligned with recent studies proposing to focus on top-ranking outlier loci in GEA analyses (e.g., 69).

**Figure 2.**
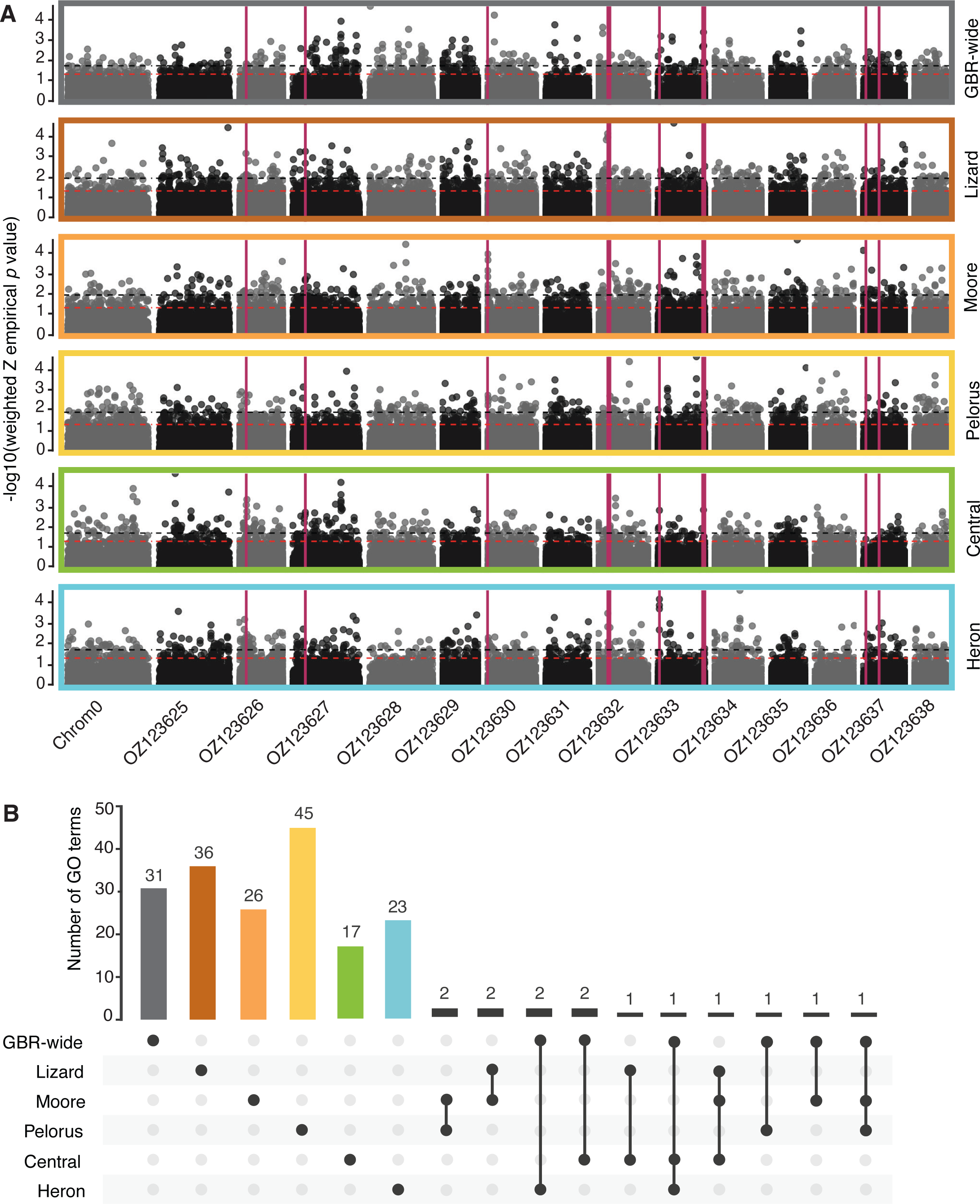
Genomic architecture and repeatability of thermal adaptation across *Stylophora pistillata* coral populations. (A) Population-level genotype-environment associations show that many 10-kb genomic windows are significantly associated with mean temperature across all chromosomes (shown in alternate colours on the x-axis). Black dotted lines show the 99^th^ percentile threshold of Weighted Z empirical *p* values and red dotted lines show the *p*=0.05 threshold. Eleven genomic windows are repeatedly associated across multiple populations (PicMin *q*<0.1, red vertical bars). From top to bottom, we show associations GBR-wide, and then for each population from North to South; (B) UpSet plot showing the overlap in enriched Gene Ontology (GO) terms (level 6) among populations. Bars in the upper panel indicate the number of enriched GO terms: population-specific terms are shown in coloured bars, while terms shared by two or more populations are shown in black bars. Dots in the lower panel represent the populations with connected dots indicating which populations share the corresponding enriched GO terms.

### Chromosomal-level visualisation of GEA analyses results

Several reference genomes exist for *S. pistillata*. Here, we selected a contig-level genome assembled from a GBR coral specimen to call SNPs because it gave us the maximum percentage of read mapping. To investigate the genomic architectures of local adaptation across chromosomes, we used RagTag v2.1.0 (70) to scaffold the contig-level assembly (GCA_032172095.1) to a chromosomal-level *S. pistillata* reference genome assembly (GCA_964205215.1) with the “scaffold” command and default parameters. We visualise the alignment between the scaffolded genome and the chromosomal genome in D-GENIES (71) and assessed the impact of scaffolding the contig-level assembly. We then projected the GEA genomic window positions onto chromosomal coordinates and the transformed data were used to generate Manhattan plots where -log_10_-transformed Weighted-Z *p*-values were plotted against chromosomal positions.

### Repeatability of thermal local adaptation across populations

To investigate whether the same genomic regions are under selection in multiple populations, we used PicMin (72). In brief, PicMin relies on order statistics to examine *p*-values (here, weighted-Z *p*-values) for the same genomic region (here, 10-kb weighted-Z windows) across multiple lineages (here, the five populations). For each population, we first transformed weighted-Z parametric *p*-values to empirical *p*-values. We then ran PicMin on windows exhibiting a weighted-Z empirical *p*<0.05 for at least one population. PicMin *p*-values were then pooled and corrected using a genome-wide false discovery rate (FDR). Windows with FDR *q*<0.1 were considered as showing strong evidence of repeatability (following (73)). This inclusive FDR-threshold of 0.1 was chosen to capture as many true positives as possible while limiting the inclusion of false positives. At a 0.1 threshold, we expect no more than 10% of significant genomic windows to be false positives. To estimate the number of expected repeated genomic windows under a model of no genotype-environment associations, we performed permutation tests (Supplementary Information).

### Functional enrichment in candidate windows under thermal adaptation

The *S. pistillata* reference genome was structurally annotated using the Egapx pipeline v.0.5.2 (https://github.com/ncbi/egapx) with 41 RNA-Seq datasets (Supplementary Information) and the taxonomic ID (50429) provided for *S. pistillata* as a guide for protein evidence. Annotations were assessed using BUSCO (74) and OMark (75) and we found high completeness and taxonomic support for the predicted genes (Supplementary Information). Predicted protein sequences from the final gene set were then functionally annotated using two complementary approaches. eggNOG-mapper (76) was run with DIAMOND search mode against the eggNOG database (released 31 March 2025), using protein input mode. To reduce taxonomically inappropriate functional transfers, searches were performed with a Eukaryota taxonomic scope and Cnidaria-restricted annotation donors (*--tax_scope eukaryota, --tax_scope_mode inner_narrowest, --target_taxa 6073*). We used sensitive DIAMOND mode with iterative searches enabled, an e-value threshold of 1e-5, a minimum alignment score of 60, minimum query and subject coverage of 50%, and non-electronic GO evidence only. Predicted proteins were additionally annotated with InterProScan v5.75-106.0 (77). InterProScan and EggNog Gene Ontology (GO) terms were then used to build a custom annotation database using the ‘AnnotationForge’ R package (97). 66,885 GO terms were identified across 15,882 unique genes, with 64% of genes having associated GO terms.

We explored signals of functional repeatability among populations by estimating Biological Processes GO term enrichment for candidate adaptive genomic regions in each population. For each population dataset, we performed GO enrichment analyses for the candidate adaptive genomic regions. We used the custom annotation database in the ‘topGO’ R package (78), with a node size of 5 and the ‘elim Fisher’ algorithm to account for the hierarchical structure of the GO graph.

Additionally, we functionally characterised loci repeatedly associated with mean temperature across populations (PicMin *q*<0.1) by intersecting the 10-kb windows with the genome annotation using bedtools intersect (79). For uncharacterised proteins, we used AlphaFold3 (80) to generate protein structure predictions and then Foldseek (81) to search for structurally homologous proteins in the available databases.

### Multivariate genotype-environment associations and predictions

To test whether redundancy in the genomic architecture of thermal adaptation (see Results) affects predictions of adaptive patterns at unsampled locations, we used a multivariate framework. We ran Redundancy Analyses (RDAs) (82) as they have been shown to be effective at identifying axes of genetic variation that covary with environmental predictors (83).

Because RDA analyses cannot handle missing data, we phased and imputed the VCF file in BEAGLE 5.3 (84, 85). To capture the environmental space, we performed a PCA on the six uncorrelated environmental variables described above, which were centred and scaled across the entire dataset. We retained the three PC axes as explanatory variables in the RDA models, which explained about 80% of the environmental variation. We performed RDAs at the GBR-wide level (i.e., all individuals) and for each population independently using the ‘vegan’ R package (86).

At the GBR-wide level, we evaluated the relative contributions of environment and geography to genomic variation, and assessed potential confounding between the two, by fitting three GBR-wide RDA models: (i) an environmental model using the first three environmental PC axes as explanatory variables, (ii) a geography model using latitude and longitude as explanatory variables, and (iii) a partial RDA conditioning on latitude and longitude to estimate the unique environmental signal independent of geographic structure. Model significance was assessed using permutation tests (999 permutations).

In addition, we evaluated the predictive ability of RDA models by using genotype-environment associations learned from a training population to predict scores for genotypes in a test population (Supplementary Information). Training datasets were either a single population or all populations combined (excluding the test population to ensure that training and test datasets were truly independent). This produced predicted genotype scores and constituted an out-of-sample prediction where the genomic data of test individuals play no role in generating their predicted scores. To quantify the predictive performance of training models, we calculated the Euclidean distance between observed (Obs) and predicted (Pred) scores across the first two RDA axes, for each individual (i):

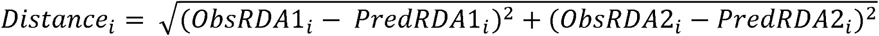

Small distances suggest high predictive accuracy while large distances suggest low predictive accuracy of genotype-environment associations from training to test population.

### Genomic offsets and predictions of population vulnerability

To predict changes in genotype-environment associations under projected climate change scenarios, and to identify potential disruptions in local adaptation, we calculated genomic offsets using the RDA framework (87). This method extrapolates contemporary genetic–environment relationships measured at sampling sites to a wider geographic area and then to future environments. The distance between contemporary and optimal future genotype scores is calculated as the genomic offset, which represent the genetic shift needed to track climate conditions (29, 34).

The models included three environmental variables: mean temperature, annual temperature range and distance to shore, as temperature is the only high-resolution environmental data available to characterise both past and predicted future climates. The two thermal metrics were selected to capture long-term and seasonal changes expected in future decades. Distance to shore was selected as a static variable to ensure that the genomic offset model does not simply recapitulate temperature changes and to capture the inshore-offshore axis of environmental variation. For the genotypes, we used all SNPs, as many empirical and simulation studies have found minimal performance differences between genomic offset models trained only on outlier loci and models trained on all available genomic markers (e.g., 35, 92–95).

This analysis requires both historical temperature data (for each sampling site, and GBR-wide) and projected future temperature data (GBR-wide). Historical and future distance to shore are the same and were calculated as described above. Historical temperature data were obtained from satellite-derived records of NOAA Coral Reef Watch, which have a 5 km resolution. We extracted these data for all reefs across the GBR, including for our sampling sites, and annual means between 1985 and 2024. For future climate data, we created an ensemble model using data between 2014 and 2100 from ten Atmospheric-Ocean General Circulation Models (AOGCMs) of the Coupled Model Intercomparison Project Phase 6 database (88): CNRM-ESM2-1, EC-Earth3-Veg, GFDL-ESM4, IPSL-CM6A-LR, MIROC6, MIROC-ES2L, MPI-ESM1-2-HR, MRI-ESM2-0, NorESM2-LM, UKESM1-0-LL. We extracted data for three Shared Socioeconomic Pathways (SSPs): SSP1-2.6, SSP2-4.5, SSP5-8.5, which represent alternative scenarios of atmospheric carbon concentration and warming trajectories (global temperature increase of ∼2.0°C, ∼2.7°C and ∼4.4°C above pre-industrial levels, respectively) (89). Daily projections were downscaled from between 80 and 500 km to 10 km using semi-dynamical shelf-sea modelling (90, 91) and we calculated decadal means. We focused our analyses on projections for two decades: 2040-2050 and 2090-2100.

First, we built a GBR-wide RDA model, as it was found to perform best in spatial genomic predictions (see Results). We did not condition the model on geography (i.e., no correction for population structure), to conservatively retain environment-associated genomic signals that co-varies with geography, since climate on the GBR is inherently spatially structured. We acknowledge this may inflate genomic offset estimates due to confounding isolation-by-distance signal (Supplementary Information). Second, we predicted present-day and future adaptive indexes for all reef area pixels across the GBR, from the historical and future (2040-2050 and 2090-2100) environmental data, respectively. The adaptive index represents the genetic similarity between these locations as a function of the environmental predictor values (87). Finally, we calculated the difference between the two predictions (historical and future) to obtain a genomic offset value, which represents the shift in adaptive index required to maintain the genotype-by-environment association that we observe today.

## Results

### Whole genome re-sequencing data

Short read whole-genome sequencing yielded a mean coverage of 9.7x. After removing five technical replicates and two low-quality samples, we obtained data for 230 individuals and 34 million SNPs. After filtering, we retained 2.4 million high-quality, bi-allelic SNPs, each with less than 20% missing data. For population structure assessment, we retained 1.3 million SNPs not in strong LD. For population-level GEA analyses, we divided the full, unfiltered dataset into six datasets based on the genomic PCA ordination results (Figure 1A), re-applied SNP filters and retained between 640,000 and 1.9 million linked SNPs.

### Environmental variation and population structure assessment

The genomic PCA based on SNPs revealed strong spatial genetic population structure (Figure 1B) with six clusters (“populations”) along the first two ordination axes: Heron Reef, Lady Musgrave Reef, Pelorus Reef, Central Reefs (Davies, Chicken and Little Broadhurst reefs, ∼10-20 km apart), Moore Reef and Lizard Reef (Figure 1A). Lady Musgrave Reef was excluded for population-level analyses due to small sample size. The environmental PCA (six environmental variables) also revealed sampling site grouping: env-PCA1 was strongly loaded by mean annual temperature (eigenvector coefficient= −0.57), monthly temperature range (−0.53) and distance to shore (0.52), separating sampling sites along a latitudinal gradient; while env-PCA2 was driven by turbidity (0.64) and light intensity (−0.54), separating shallow and deep sites within each reef (Figure 1C). Thus, the six populations inhabit distinct environments broadly (env-PCA1), with depth-associated gradients replicated within each (env-PCA2).

### Genomic architecture of thermal local adaptation across populations

We first focused on mean temperature, given its relevance for adaptation under pervasive warming (Supplementary Information). Mean temperature correlates with geographic distance as captured by latitude (Figure 1A, env-PC1) and therefore false positives arising from differentiation by distance could affect GEA analyses focusing on range-wide thermal adaptation. Here, however, we tested for GEAs within each population independently, decoupling temperature from geography (formally addressed in the next subsection). Permutated weighted-Z scores were centred around zero, while observed scores showed broader distributions, indicating an excess of higher absolute values compared to the null (Figure S3). At an empirical *p*-value threshold of 0.05, we identified between 814 and 1,359 candidate adaptive genomic windows per population, suggesting a polygenetic basis for thermal adaptation.

Using PicMin (72) we identified eleven genomic windows significantly associated with temperature across populations, at False Discovery Rate (FDR) *q*<0.1 (one significant in three populations, three in four populations, and seven in all five; Figure 2A, Figure S4), spanning six RagTag chromosomes. At FDR *q*<0.5, 188 genomic windows were significant. Results were consistent using 20-kb windows, with four repeated windows across all populations (*q*<0.1, Figure S5). Under a model of no repeatability using permutations, a mean of zero genomic windows were significant for PicMin at *q*<0.1.

### Functional annotations of candidate adaptive loci

To assess repeatability at the functional level, we conducted GO term enrichment analyses on candidate genomic windows under thermal selection within populations (WZA empirical *p*-value >99th percentile; 319 windows per population). Enriched GO terms were related to processes including transport, gene regulation and regulatory stress response (Figure S6, Figure S7), and several of these terms are consistent with known mechanisms of thermal adaptation in corals, including response to temperature stimulus and protein refolding. GO enrichment was largely population-specific: only 7% (14/192) of enriched terms were shared by two or three populations (Figure 2B).

The eleven genomic windows repeatedly associated with temperature (PicMin *q*<0.1) overlapped with15 genes, including 13 protein-coding genes and 2 non-coding RNAs (Table S2), that play roles in micronutrient acquisition and cellular metabolism, protein turnover, apoptotic signalling, and cell adhesion and recognition.

### Predictability of local adaptation across the species’ range

Given low repeatability in the genomic basis of thermal adaptation, we assessed how this influences the predictability of genome-wide local adaptation patterns. RDA models were built for all populations together (i.e., GBR-wide) as well as for each of the five populations independently (Figure S8) and all models were significant (permutation testing, *p*<0.05, Table S3). GBR-wide variance partitioning showed that environmental and geographic predictors each explained a small but meaningful proportion of genomic variance (3.0% and 2.8% adjusted R², respectively). After conditioning on geography (partial RDA), the unique environmental signal was reduced to 1.4%, as 38% of the full model’s explainable variance was confounded with geography (Figure S9). Of this partial model’s explainable variation, 45% was captured by env-PC1 (shore distance and thermal metrics).

RDA models were then used in a predictive framework, using genotype-environment relationships learned from one geographic region (training dataset) to predict genotype scores for another geographic region (test dataset). RDA models trained on a single reef population resulted in larger Euclidean distances between predicted and observed genotype scores (Figure 3, Figure S10), indicating poor ability to predict putatively adaptive genomic patterns, likely because they were fit to a narrow range of environmental variation, typically outside of that experienced in the test population. In contrast, the models trained on the GBR-wide environmental range (except the test population) predicted genotype scores that closely matched observed values (Figure 3, Figure S10), consistent with interpolation within a broader environmental space. Among the three GBR-wide models, the partial RDA model (conditioning on geography) consistently had the greatest prediction accuracy across test populations.

**Figure 3.**
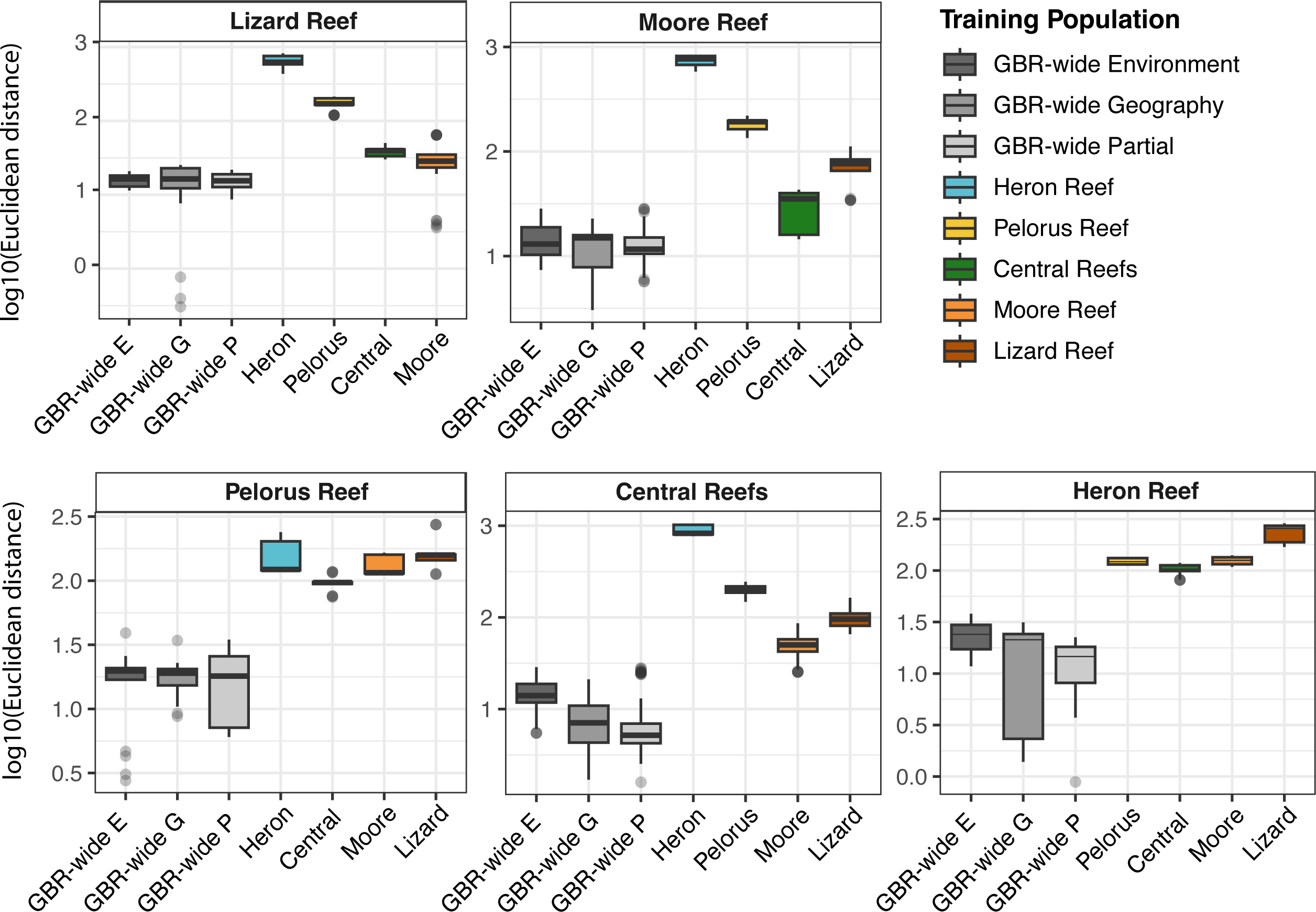
Range-wide models have higher performance in predicting genotype-environment associations across *Stylophora pistillata* populations. Each panel focuses on a specific test population and shows the distribution (boxplot) of the log10-transformed Euclidean distances between observed and predicted genotype RDA scores, calculated from the two first Redundancy Analysis axes. Predicted scores were obtained from RDA models trained on GBR-wide genotype-environment associations (“GBR-wide E” boxplots), GBR-wide genotype-geography associations (“GBR-wide G” boxplots), GBR-wide genotype-environment associations conditioned on geography (“GBR-wide P” boxplots) and on single-population genotype-environment associations (coloured boxplots). Lower distances correspond to better predictive accuracy. For example, the first panel shows that the models trained on GBR-wide genotype-environment associations have the highest performance (i.e., lowest Euclidean distances) in predicting genotype scores at Lizard Reef, whilst the model trained on genotype-environment associations at Heron Reef has the lowest performance. Note: y-axis scale varies among panels.

### Genomic predictions of local (mal)adaptation under climate change

Rising ocean temperatures directly threaten coral populations. To identify *S. pistillata* GBR populations at higher risk of future maladaptation, we calculated genomic offsets (following 87). Larger values suggest that a greater genomic shift will be required to maintain present-day genotype-by-environment association (i.e., greater maladaptation) (29, 34). Genomic offsets varied both among and within geographic regions (Figure 4). For the 2040-2050 decade, all three climate change scenarios produced similar genomic offset distributions, with higher values at northern and inshore reefs (Figure 4). Projections for 2090–2100 diverged more strongly across scenarios: offsets increased significantly at northern reefs under SSP2-4.5 relative to SSP1-2.6, and further increased range-wide under SSP5-8.5 (Figure 4). These later projections included temperature values exceeding the range-wide historical maximum and generated genomic offsets for novel thermal environments, reaching +2.3°C above historical maximum in the Far North GBR under SSP5-8.5 in 2090-2100 (Figure S11).

**Figure 4.**
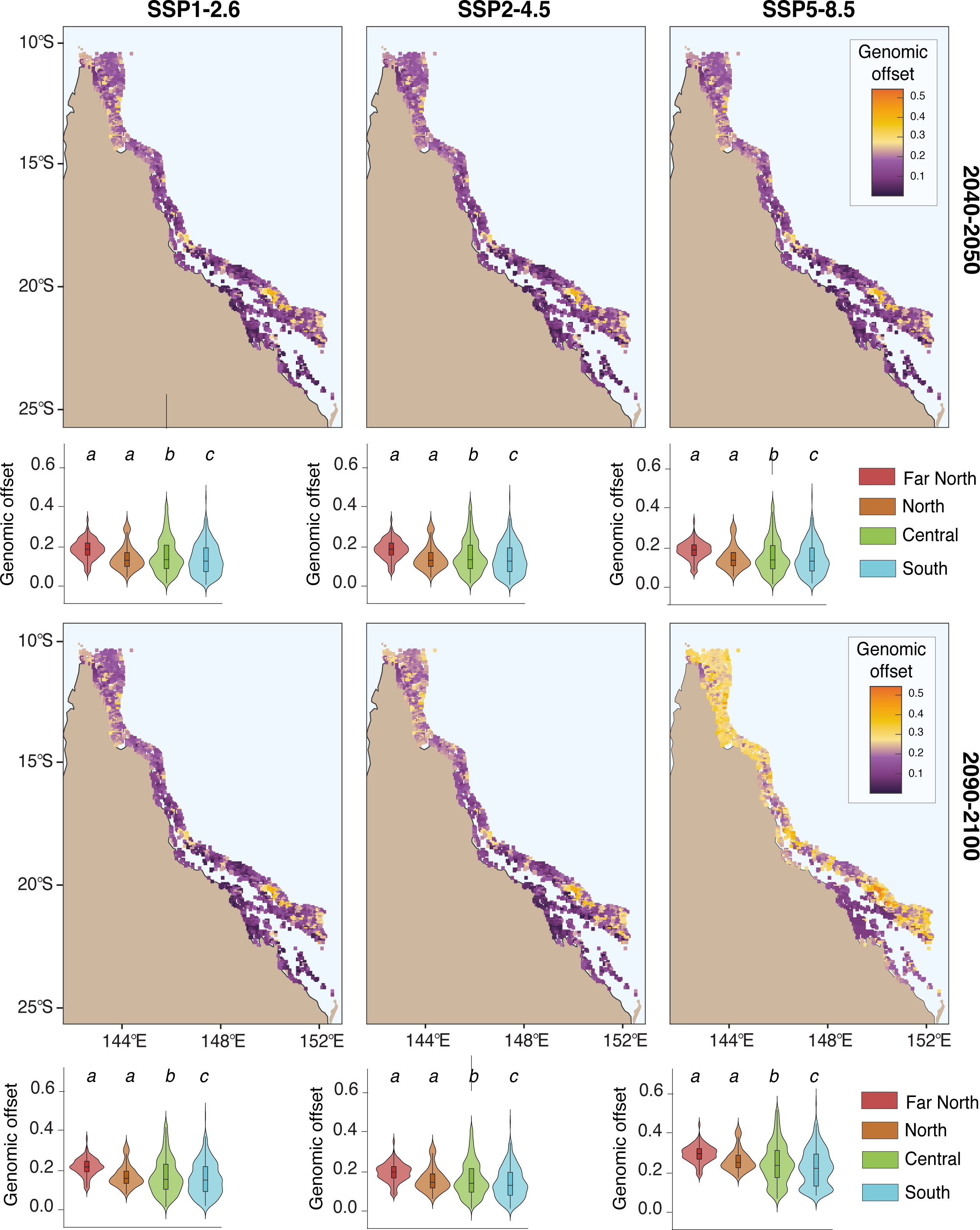
Spatial variability in genomic predictions of local adaptation under climate change across Great Barrier Reef *Stylophora pistillata* coral populations. Distribution of genomic offset values across the Great Barrier Reef under three climate change scenarios projected for two decades (2040-2050, top panels; and 2090-2100, bottom panels). SSP1-2.6 represents a low greenhouse gas emission scenario, SSP2-4.5 an intermediate emission scenario and SSP5-8.5 a high emission scenario. Future climate data was obtained using an ensemble of ten CMIP6 models. Higher values indicate that a greater genomic shift is required to maintain present-day genotype-by-environment association. Summary of genomic offset distribution across four latitudinal regions are shown for each scenario-decade as violin plots, for the Far North (> 15.5°S), North (−18°S to −15.5°S), Central (−21.5°S to −18°S) and South (< −21.5°S) regions. Boxplots inside the violin plots highlight the median and interquartile range. Different letters above the violin plots represent groups that are significantly different based on Tukey’s post-hoc tests (*p*<0.05).

## Discussion

Local adaptation will be critical for shaping a species’ response under changing environmental conditions (12, 13, 33), yet we know little about its genetic basis across distinct populations. In this study, we show that genomic signatures underlying thermal adaptation are geographically restricted for the short-dispersal coral *Stylophora pistillata* (Taxon1; sensu 55). Despite limited repeatability at the locus level, geographically extensive genotype-environment associations accurately predicted genome-wide adaptive patterns across populations and we detected high regional variability in genomic offsets under future climate scenarios across the Great Barrier Reef. These results highlight the importance of spatial scales in detecting signatures of environmental selection and refine the applicability of genomic models for forecasting local adaptation.

By leveraging strong genetic differentiation among *S. pistillata* GBR populations (Figure 1B) and environmental heterogeneity within reefs (Figure 1C), we found that most candidate genomic regions under thermal selection in one population were not candidates in others. This aligns with theoretical expectations for polygenic climate adaptation (96) and prior reports of non-parallel local adaptation among populations and closely related species (e.g., 97, 98). Such genomic redundancy may reflect historical allele frequency differences among these populations. Alleles at intermediate frequencies are more likely to contribute to evolutionary responses (99), such that loci common in one population but rare in others are unlikely to repeatedly contribute to adaptive responses. Population differentiation in allele frequencies due to demography and drift in *S. pistillata* are implicated by previous estimates of low genetic diversity (π∼0.01), small effective population sizes (50<*N_e_*<960) and strong genetic differentiation among populations (0.04<F_ST_<0.15) due to limited gene flow (*N_e_*m<4 migrants/generation) (54). Future work could test the contribution of locus-specific frequency variation by examining the frequency distributions of the identified candidate loci.

Despite high genomic redundancy, we identified eleven genomic windows, overlapping with thirteen genes, that were repeatedly associated with temperature across two or more populations (Figure 2A, Figure S4). By looking at shared signals across populations, this analysis bypasses the known challenge of high false positive rates in broad-scale GEA analyses when environmental variation covaries with neutral population genetic structure (30–32). Because genetic drift is unlikely to produce parallel signatures, repeatability provides compelling evidence for the adaptive role of these genomic regions. Among the proteins encoded in these genomic regions, peroxidasins are frequently differentially expressed in coral heat stress studies (e.g., 100, 101) and sulfotransferase family proteins are putatively under strong positive selection between depth-segregated coral lineages (102). Given the evolutionarily recent population divergence (∼400,000 generations ago, 54), these adaptation signals likely represent shared ancestral genetic variation, rather than independent repeated mutations. As small-effect adaptive loci are difficult to detect in genome scans (25, 72), we might not have identified all repeatedly selected upon loci, but the thirteen genes identified here warrant further examination.

Because different large-effect loci can contribute to the same biological function, functional-level repeatability might be expected even in the absence of locus-level repeatability (22). However, candidate adaptive genomic regions in distinct populations were enriched in mostly different biological processes (Figure 2B). This suggests that thermal adaptation can be reached via different functional routes and aligns with the hypothesis that highly complex traits might not show functional convergence (22). Alternatively, findings of low repeatability at both the genomic and functional levels could also reflect differences in the adaptive landscape among populations. All analysed populations experience comparable levels of thermal variability across habitats, but under vastly different ranges of absolute temperatures (Figure S11), which may require different physiological, and thus genetic, adaptive solutions.

Complementing genome scan results, we asked whether geographically restricted adaptive architectures constrain the predictability of adaptive genomic compositions across the species’ range. RDA models trained on range-wide data predicted genotype scores at a specific location with high accuracy (Figure 3), indicating that these models capture reliable genome-wide isolation-by-environment patterns, even if they cannot accurately identify causal loci (which vary across populations). In an RDA, each locus is first modelled individually as a function of the environmental variables such that minor differences in allele frequency among individuals or populations do not prevent the detection of shared environmental associations. The aggregate fitted values across all loci can therefore still reflect the same overall pattern, even if some loci do not show statistically significant associations on their own. These observations echo previous simulation-based results that recommended the use of ordination methods for predicting individual environmentally adaptive traits instead of identifying specific outlier loci (32).

In contrast, RDA models trained on a single reef population performed poorly (Figure 3), likely reflecting different genotype–environment relationships among populations and greater uncertainty when extrapolating elsewhere in the range. Notably, predictive performance varied among training populations, with models trained on Heron Reef performing the worst (Figure 3), possibly because of the unique environmental conditions (Figure 1C) associated with the Capricorn Eddy (103). Conversely, models trained on the Central Reefs population had generally higher predictive performance, which can be attributed to the central position of this population along the first axis of the environmental space that explains most variation of the GBR-wide RDA model (Table S3). Overall, the more geographically and environmentally distant the training population was from the test population, the less accurate the predictions. This reinforces the importance of environmental breath in training data and model specificity in landscape genomics (35, 104–106).

Predicting population (mal)adaptation to climate change using genomic offsets is increasingly popular (107) and relies on projecting GEAs across a wide geographical range under present-day and future climatic conditions. Having shown that range-wide genotype-environment patterns can be extrapolated reliably across space, we hypothesised that extrapolation forward in time might also be robust, provided that future climates are represented in the training dataset. We show that predicted (mal)adaptation is highly variable across GBR *S. pistillata* populations (Figure 4). In the near future (2040–2050) and under low (SSP1-2.6) and intermediate (SSP2-4.5) carbon emission scenarios, the highest genomic offsets (i.e., least adapted to future conditions) occurred mostly in the northern GBR, while the lowest genomic offsets (i.e., most adapted to future conditions) were patchily distributed across the central and southern regions (Figure 4). Our findings complement previous modelling studies identifying possible thermal refugia (91, 108, 109), but indicate that thermal and genetic refugia might not always coincide. Under intense warming (SSP5-8.5) in 2090-2100, this spatial heterogeneity disappears, with all reef populations showing similarly high disruptions of local adaptation (Figure 4). Importantly, population adaptation to climate change will not only depend on the disruption of genotype-environment relationships but also on other processes shaping adaptive genetic variation (e.g., effective population size, migration), which are not considered here.

Nonetheless, genomic offset models remain useful to identify populations with different genotype-environment matching to future conditions — assuming no adaptive gene flow. For long-lived, sessile and short-dispersal species such as *S. pistillata*, this assumption is plausible in the short term. One important limitation, however, is the extent to which genomic offset models might be affected by environmental novelty. In this study, we show that accurate prediction of local adaptation highly depends on the range of environmental variation captured by the training dataset. As terrestrial (110) and marine (111) organisms are expected to experience environmental change exceeding three standard deviations relative to historical means, genomic offset estimates are likely to be inaccurate in regions projected to change above recent historical conditions. In our models, this is the case for the northern GBR under SSP5-8.5 in 2090-2100, where mean annual temperature exceeds 30°C compared to a range-wide historical maximum of 27.7°C (Figure S11).

To conclude, we show that genome-wide signals, rather than locus-specific signatures, can drive reliable genomic forecasting. Thus, a mechanistic understanding of climate adaptation might not be necessary to inform conservation actions and efforts might be better spent characterising genotypes across a broad range of environments. Here, we identified coral populations that may be at higher maladaptation risk in the coming decades, which could be prioritised in reef restoration interventions. For example, a reverse genomic offset metric could help identify suitable source populations for strategies such as assisted gene flow (7). While promising, experimental validation of these correlative models is crucial for assessing their accuracy. For corals, common garden and reciprocal transplant experiments can be challenging, given the species’ long lifespans, high costs, and regulatory restrictions for translocations across large distances. Instead, observational data, collected before and after natural marine heatwaves for example, may offer a suitable validation alternative.

## Acknowledgments

The coral samples in this study were collected from the traditional sea Country of the Bailai, Gurang, Gooreng Gooreng, Taribelang Bunda, Bindal, Manbarra, Gunggandji, Nguurruumungu, Dingaal and Thanil peoples with their Free Prior and Informed Consent. The authors are grateful for the granting of consent and we pay our respects to Elders past, present and emerging. We thank the ecoRRAP team for organising field expeditions, S. Howitt and I. Byrne for their assistance with laboratory work and sample collection, P. A. Gagnaire and O. Gaggiotti for providing valuable feedback on an earlier draft. This work was supported by the Reef Restoration and Adaptation Program, funded by the partnership between the Australian Government’s Reef Trust and the Great Barrier Reef Foundation and by an Australian Government Research Training Program (RTP) Scholarship.

## Data Availability

Raw sequencing data are deposited on NCBI (BioProject PRJNA1404865); metadata for all samples is deposited on GEOME (https://n2t.net/ark:/21547/Gkw2); scripts to reproduce analyses and figures are available on GitHub (https://github.com/zoemeziere/GEA_Stylophora).

## Notes

### Competing Interest Statement

The authors have declared no competing interest.

### Summary of Updates

New format, with results after methods. Updated Acknowledgments section.

