## Supplementary Information for "Genomic repeatability and predictability of climate (mal)adaptation in a reef-building coral"

**Supplementary methods and extended explanations of data analysis decisions**

**Using mean temperature to test genomic signatures of local adaptation**

We focused on mean temperature to investigate genomic architectures of local adaptation within populations for several reasons. First, temperature is one of the most important environmental components to coral physiology and survival: it is central to regulating the coral host’s metabolism, symbiont performance and it also sets the threshold for coral bleaching (1, 2). Second, thermal heterogeneity within reefs is well documented (e.g., (3)), with meaningful temperature differences among reef habitats and along the depth gradient, which is the spatial scale at which *S. pistillata* disperses and recruits (4). Temperature variation within regional population therefore provides a tractable selective gradient, independent of the broad north-south temperature gradient that co-varies with neutral genetic structure among populations. Third, using a single, ecologically meaningful variable allowed us to directly compare the genomic architecture underlying the same selective pressure in the five *S. pistillata* populations, which is a necessary condition for testing repeatability of adaptive patterns. Including additional environmental variables into this analysis would have complicated the attribution of shared genomic signals to a common selective agent. Finally, future changes in mean temperature are the most threatening impact of climate change for coral populations, making it the most relevant variable in adaptive capacity forecasting.

We recognise that using modelled temperature data (e.g., eReefs GBR1 product) is an imperfect proxy for the selective environment experienced by the coral colonies on the reef. Although we used the highest resolution products available for the Great Barrier Reef, they might be too coarse to capture the fine-scale thermal heterogeneity that characterises reef environments. Notably, however, a recent comparison of satellite sea surface temperature products for a Pacific coral reef system found that a lower-resolution dataset outperformed a higher-resolution alternative at predicting *in situ* temperatures and past bleaching severity, likely due to differences in underlying data density and processing protocols rather than resolution alone (5). Ideally, *in situ* temperature loggers deployed for long time periods across all our sampling sites would provide a more accurate representation of local thermal conditions. However, this was not feasible for this study and it is a known challenge in empirical seascape genomic studies. It is therefore possible that the true strength of thermal selection within populations is greater than we detected.

**Kendall’s τ without population structure correction in genotype-environment associations**

Many methods have been developed to find associations between genomic and environmental data. Here, we calculated Kendall τ correlation coefficients between sampling site-level allele frequencies and mean temperature data. Unlike other GEA methods, Kendall τ is a rank-based correlation method that can handle ties in the data and is non-parametric (i.e., does not assume any particular distribution of allele frequencies or environmental variation).

We also favoured this simple approach over methods that attempt to correct for spatial autocorrelation between population structure environmental variation. Correcting for population structure in a GEA analysis limits the number of false positives but also reduces power to detect true positives. Conversely, approaches that do not correct for population structure have higher power to detect true positives at the cost of more false positives (6, 7). This is particularly a challenge when axes of environmental variation align with axes of population structure (8, 9), as is often the case in for wide-ranging species, and is the case for mean temperature and geography at the whole GBR scale. In our study, we decided to take the more inclusive approach of not correcting for population structure, in order to capture as many true signals for each population. We accepted an unknown number of false positives because we were interested in the repeatability of association signals across populations, where it is unlikely that the same false positives would manifest across populations. This framework therefore addresses the challenge of collinearity between neutral population structure and environmental variation.

**The Weighted-Z Analysis approach and genomic window sizes**

The Weighted-Z Analysis approach leverages signals of elevated linkage among nearby SNPs that share the same evolutionary history and this approach has similar or improved power as well as lower error rates compared to other GEA methods (7). In window-based genome scans, the window size is chosen such that all SNPs on the same window have the same coalescent history and reflect the same genealogy. This is because SNPs in tight linkage will recombine less frequently than unlinked sites. Since we do not have recombination rate estimates for *S. pistillata*, we examined LD decay by calculating squared inter-variant allele count correlations (R²) in PLINK (10) and estimated the distance over which recombination breaks down associations among SNPs (R²<0.1-0.2). Booker et al. (7) recommend choosing a window size that is wider than the LD decay distance. We found that LD decays (R²<0.2) at about 5kb (Figure S12) and therefore used windows of 10kb. This choice of window size is also motivated by the distribution of gene lengths (IQR: 1.8-8.5kb) (Figure S13).

**Permutation testing for the Weighted-Z Analysis**

For each population, we shuffled the environment among sampling sites 1,000 times to create a null of no genotype-environment associations and re-computed weighted-Z scores on the permuted datasets. For each window, the observed weighted-Z score was compared to the null distribution generated from all permutations and we calculated an empirical *p*-value as the proportion of permuted weighted-Z scores greater than or equal to the observed value.

**Using RDAs to predict genotype scores under known genotype-environment associations**

RDA scores are individual-level projections onto the constrained axes – that is, coordinates in the ordination space defined by the environment-genotype relationships. To generate these scores, the RDA regresses each SNP genotype against the environmental predictors across all training individuals. This produces a set of regression coefficients that encode how each locus covaries with each environmental predictor. Using these training regression coefficients, we built predictive RDA models and applied them to sets of test populations; effectively generating predicted genotype scores for the test individuals. This was done using the *predict()* function from the ‘vegan’ R package with *type = "lc"*, and the environmental values for the test individuals. Thus, by comparing the observed genotype score and predicted genotype score for a given individual, we can estimate where an individual's genome places them in ordination space versus where their environment alone predicts they should be.

**RDA model selection for genomic offset analyses**

For projecting genotype-environment associations into the future and estimate genomic offsets, we used an RDA model without conditioning on geography. This decision was made to conservatively retain as much environment-associated genomic signal as possible, including from the environmental variance that co-varies with geographic position. While we cannot fully disentangle environment-driven (i.e., local adaptation) and geography-driven (i.e., isolation by distance) genomic variation, climate variation is inherently spatial. On the GBR, environmental variation (including in temperature) is structured along a latitudinal axis. This means that the proportion of environmental variation that is confounded by geography might still be important for explaining coral spatial adaptation patterns and predicting (mal)adaptation patterns under future climatic conditions. We acknowledge that neutral isolation by distance signal may inflate genomic offset estimates from a model that is not conditioned on geography.

**Supplementary Results: Reference genome annotation and quality assessment**

The final egapx genome annotation comprised 30,530 genes, 28,285 mRNAs, 2,432 non-coding RNAs, 5,880 pseudo transcripts and 28,285 CDSs. Of the 30,530 annotated genes, 23,028 (75.4%) were protein-coding. Of the 28,285 mRNA models, 83.8% (23,692) were fully supported by transcript evidence.

Annotation completeness and taxonomic consistency were evaluated using OMark with **Eumetazoa** as the reference clade. Of 3,244 conserved hierarchical orthologous groups (HOGs), 89.5% (2,902) were recovered as single-copy genes. A total of 181 HOGs (5.6%) were missing, indicating high overall completeness of the gene set. Of the 23,028 predicted proteins, 67.5% (15,543) were assigned to gene families consistent with Eumetazoa. No proteins were identified as contaminants and 28.6% (6,575) could not be confidently assigned to known gene families and were classified as unknown. Eumetazoa was identified as the sole dominant clade, with 71.5% of proteins mapping to this clade, providing further support for the taxonomic coherence of the annotation.

BUSCO analysis (metazoa_odb10) showed high completeness, with 94.7% of genes recovered as complete. The majority of complete BUSCOs were single copy (94.1%).

**Supplementary Figures and Tables**

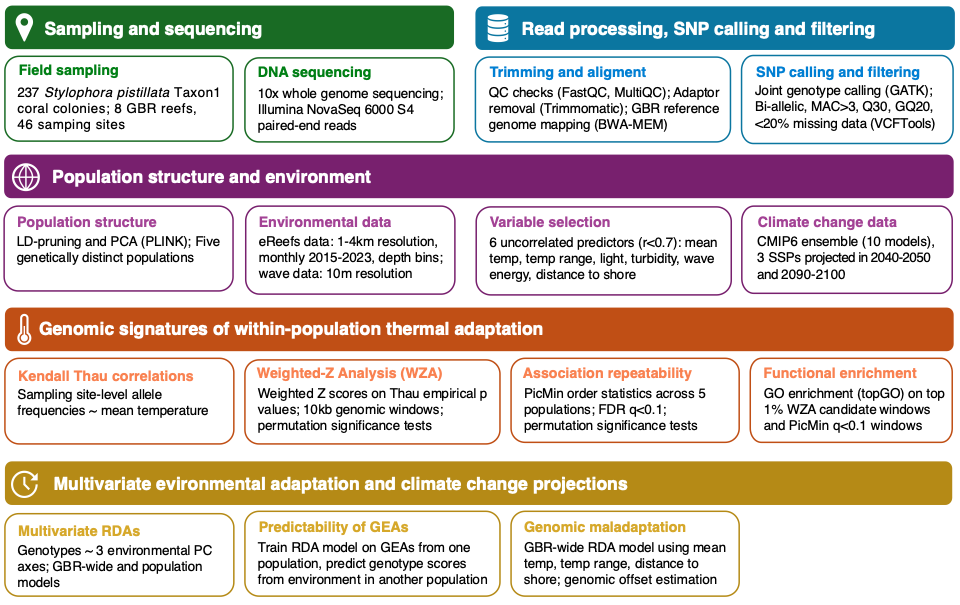

**Figure S1. Schematic overview of the study design and analytical workflow used to examine climate adaptation in *Stylophora pistillata*.** We include details on sample collection, whole-genome sequencing, SNP calling and filtering, population structure and environmental data acquisition, detection of genomic signatures of thermal adaptation and their repeatability across populations, functional enrichment analyses, multivariate genotype–environment predictive modelling and estimation of genomic offset under future climate scenarios.

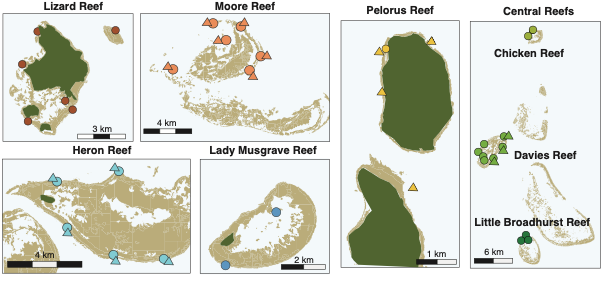

**Figure S2. Maps showing the sampling sites at each reef.** *Stylophora pistillata* coral colonies were sampled across different habitats and at shallow (4-7 meters, circles) and deeper sites (12-15 meters, triangles). This nested sampling design allowed us to sample coral colonies inhabiting environmental diverse locations.

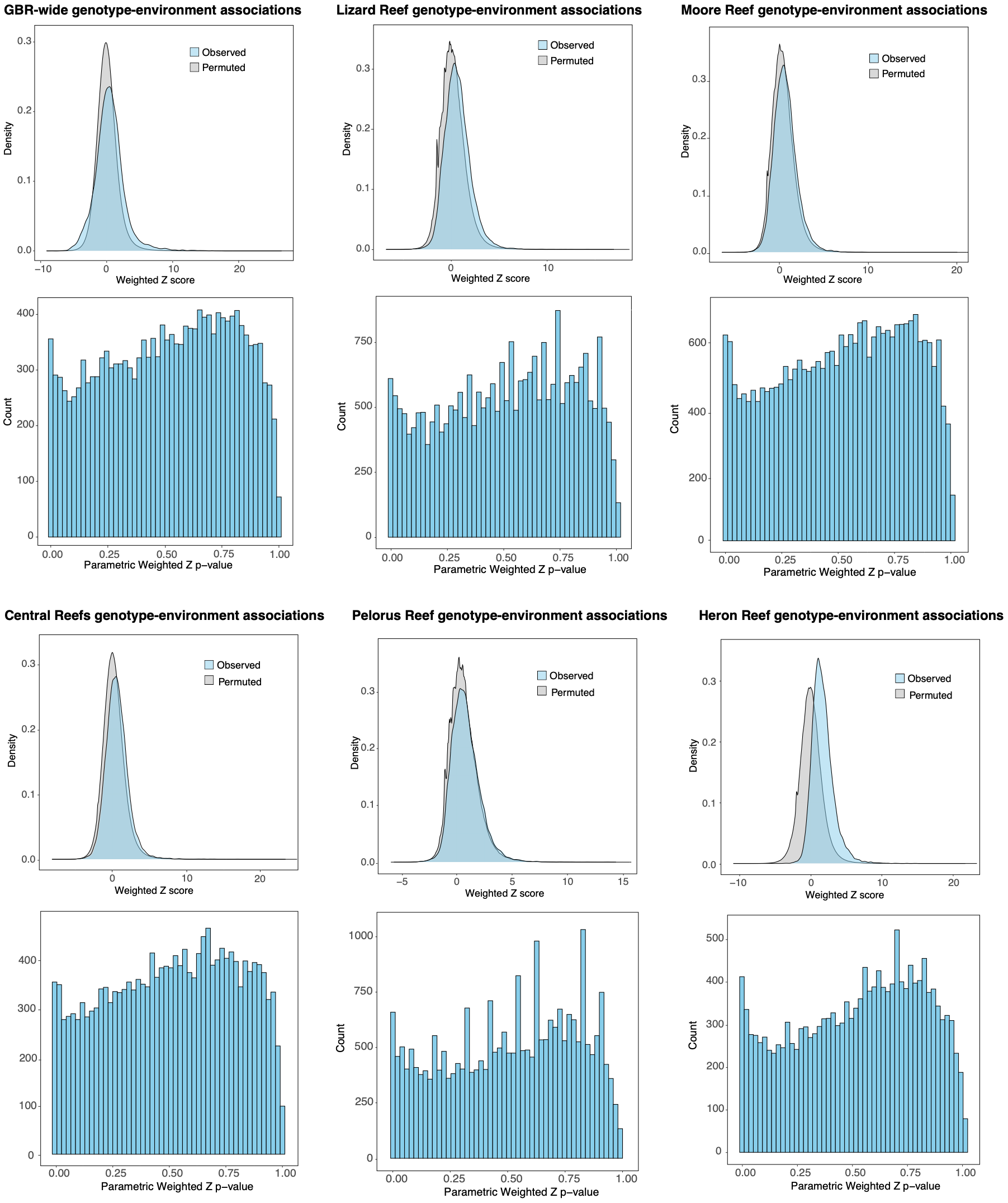

**Figure S3. Distribution of observed and permutation-based Weighted-Z Analysis (WZA) scores and corresponding empirical *p*-values.** For each population, the top panel shows the distribution of observed (blue) and permuted (grey) WZA *Z*-scores, where the broader distribution for the observed scores compared to the permuted scores indicates an excess of stronger genotype-environment associations in the empirical data. For each population, the bottom panel shows the distribution of parametric Weighted-Z *p*-values.

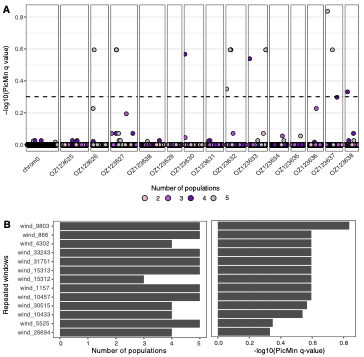

**Figure S4. PicMin analyses reveal eleven genomic windows repeatedly associated with mean temperature variation across *S. pistillata* populations.** A) Manhattan plot showing the strength of temperature-association repeatability across populations. The dashed line indicates the genome-wide significance threshold of *q* = 0.1; B) Number of populations in which these windows are significant (PicMin *q*<0.1) (left panel) and statistical support for repeatability as -log10 transformed PicMin *q* value (right panel).

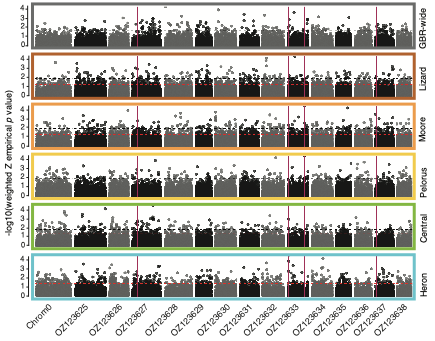

**Figure S5.** **Signatures of within-reef thermal adaptation and repeatability across populations using 20-kb windows to calculate weighted-Z scores.** Population-level genotype-environment associations show many windows significantly associated with mean temperature across chromosomes (shown in alternate colours on the x-axis). Black dotted lines show the threshold for the top 1% genomic windows with lowest Weighted Z empirical *p* value and red dotted lines show the *p*=0.05 threshold. Four genomic windows are repeatedly associated across multiple populations (PicMin *q*<0.1, red vertical bars).

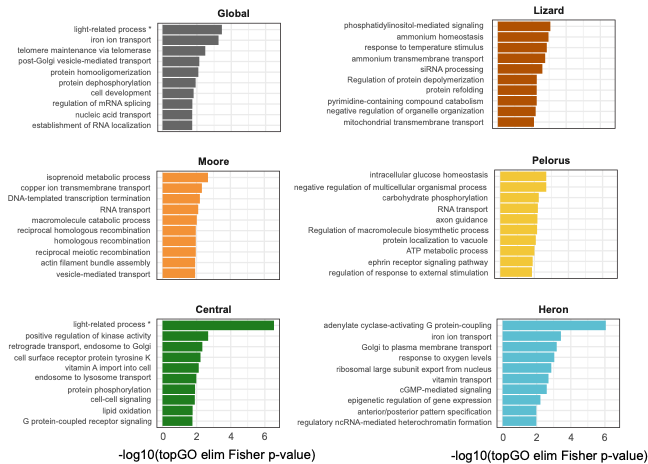

**Figure S6. Most significantly enriched Gene Ontology (GO) terms for each population**. The top ten level 6 Gene Ontology (GO) terms most significantly enriched in genomic windows identified as candidate genomic regions under thermal adaptation (top 5% WZA empirical *p*-value), for each population. * indicates a raw GO term that was updated to a more general process.

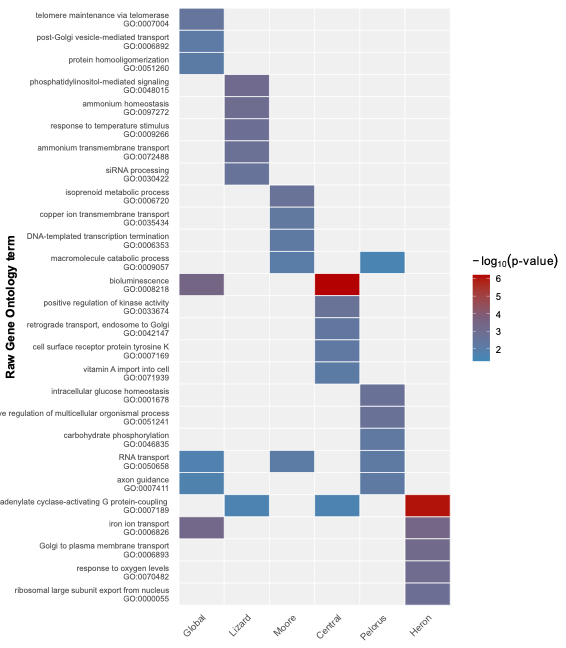

**Figure S7. Repeatability of significantly enriched Gene Ontology (GO) terms**. GO terms most significantly enriched in genomic windows identified as candidate genomic regions under thermal adaptation (top 5% WZA empirical *p*-value) (rows) for each population (columns). The cell colour indicates the significance (empirical *p*-value) of the GO term enrichment. GO terms are ordered to highlight population-specific enrichment patterns.

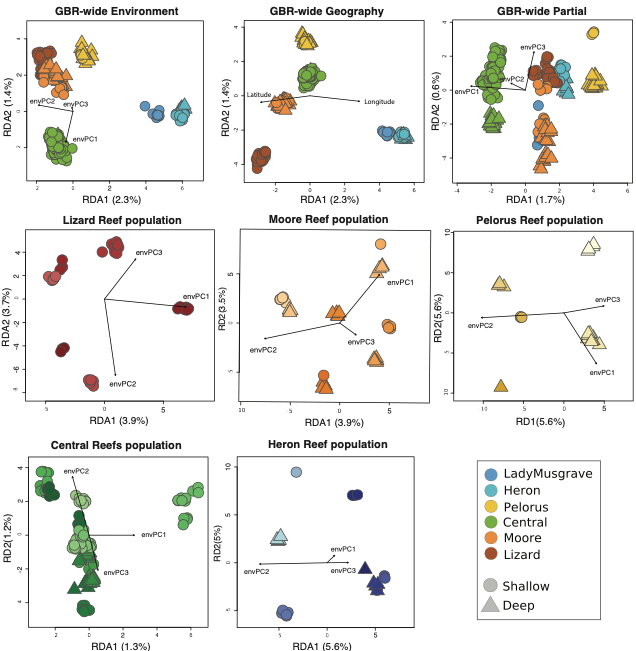

**Figure S8. Biplots of RDA models used to predict environmental adaptation.** GBR-wide and population-level RDA models are presented. Three GBR-wide models were evaluated: with environmental data, with geographic data, and with environmental data conditioned on geographic data. For the environmental data, six environmental variables (mean temperature, temperature annual range, mean light intensity, mean current velocity, mean turbidity and distance to shore) were reduced in dimensionality using PCA. The first three environmental PC axes (envPC1, envPC2 and envPC3) which were used to constrain the environmental RDA models. Samples are coloured and shaped using the same codes used in Figure 1. Axis percentages represent the proportion of genomic variance explained by the environmental predictors captured by each canonical axis.

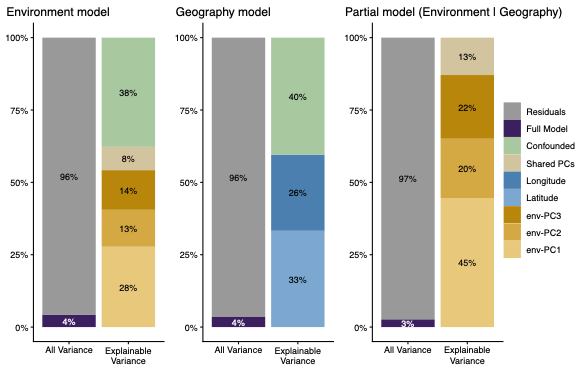

**Figure S9**. **Variance partitioning of genomic variation in Stylophora pistillata coral genotypes across the Great Barrier Reef.** Each panel shows the proportion of total genomic variance (left bar, "All Variance") and the proportion of explainable variance (right bar, "Explainable Variance") attributed to different predictors. The RDA environment model (left panel) includes three principal components of six environmental variables, explaining 4% of total genomic variance. The geography model (centre panel) includes latitude and longitude, also explaining 4% of total variance. The partial model (right panel) shows the unique environmental contribution after conditioning on geography, explaining 3% of total genomic variance. Within the explainable variance bars, coloured slices indicate unique contributions of individual predictors (PC1–3, latitude, longitude), the confounded fraction represents variance shared between environment and geography that cannot be uniquely attributed to either, and the shared PCs fraction represents variance jointly explained by multiple environmental PCs.

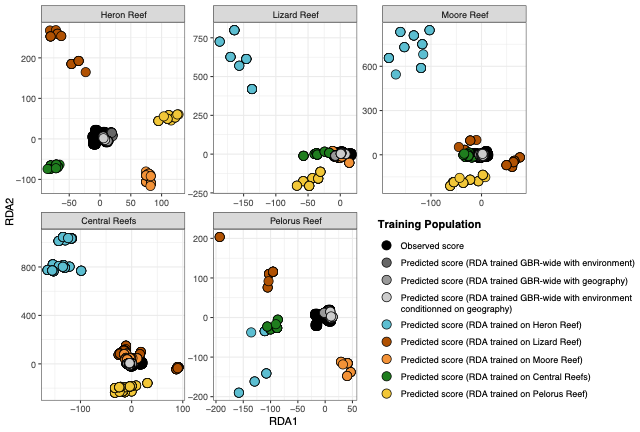

**Figure S10. Observed and predicted genotype scores from RDA models.** Each panel is a test population where genotype scores are being predicted, and the dotes represent genotype RDA scores. Black dots are the observed genotype scores in each population, and coloured dots are the predicted genotype scores from models trained on different training populations (colours indicate the training population).

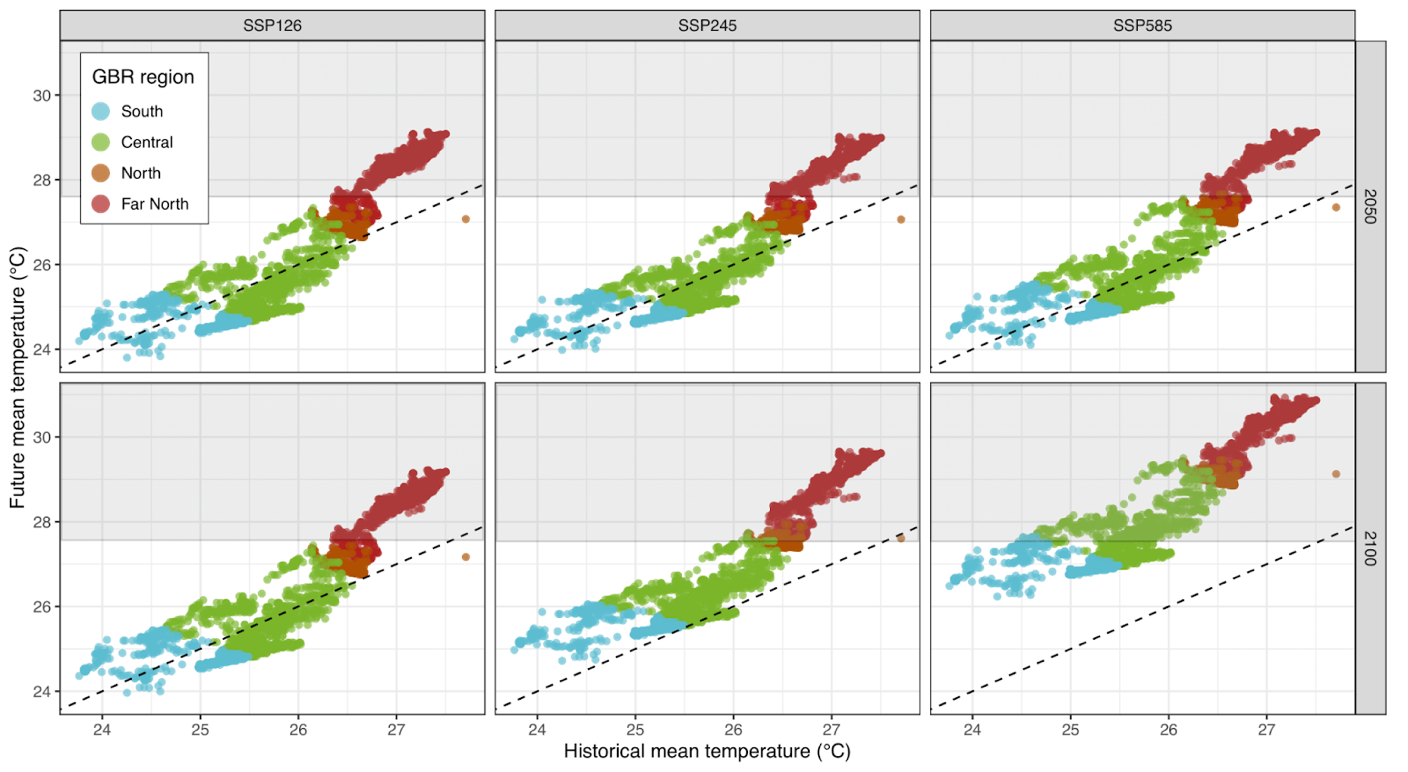

**Figure S11. The extent of future environmental novelty differs across the Great Barrier Reef and among climate change scenarios.** Plots show historical mean temperature and predicted mean temperature for all spatial data points, under three climate change scenarios and two decades. Shaded grey boxes delineate areas of environmental novelty, where predicted future mean temperature exceeds maximum historical temperature (27.7°C). Data points are coloured by geographic region.

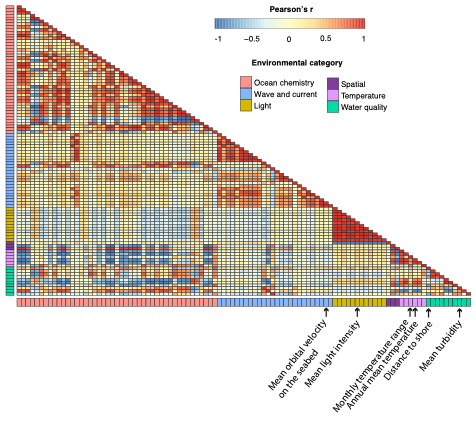

**Figure S12.** **Heatmap showing correlation coefficients (Pearson’s r) among all environmental variables considered prior to variable reduction.** Variables are grouped and color-coded by category to highlight related environmental factors for better readability. Variables used to build models in this study are denoted with arrows. The full dataset is available as an attached supplementary file.

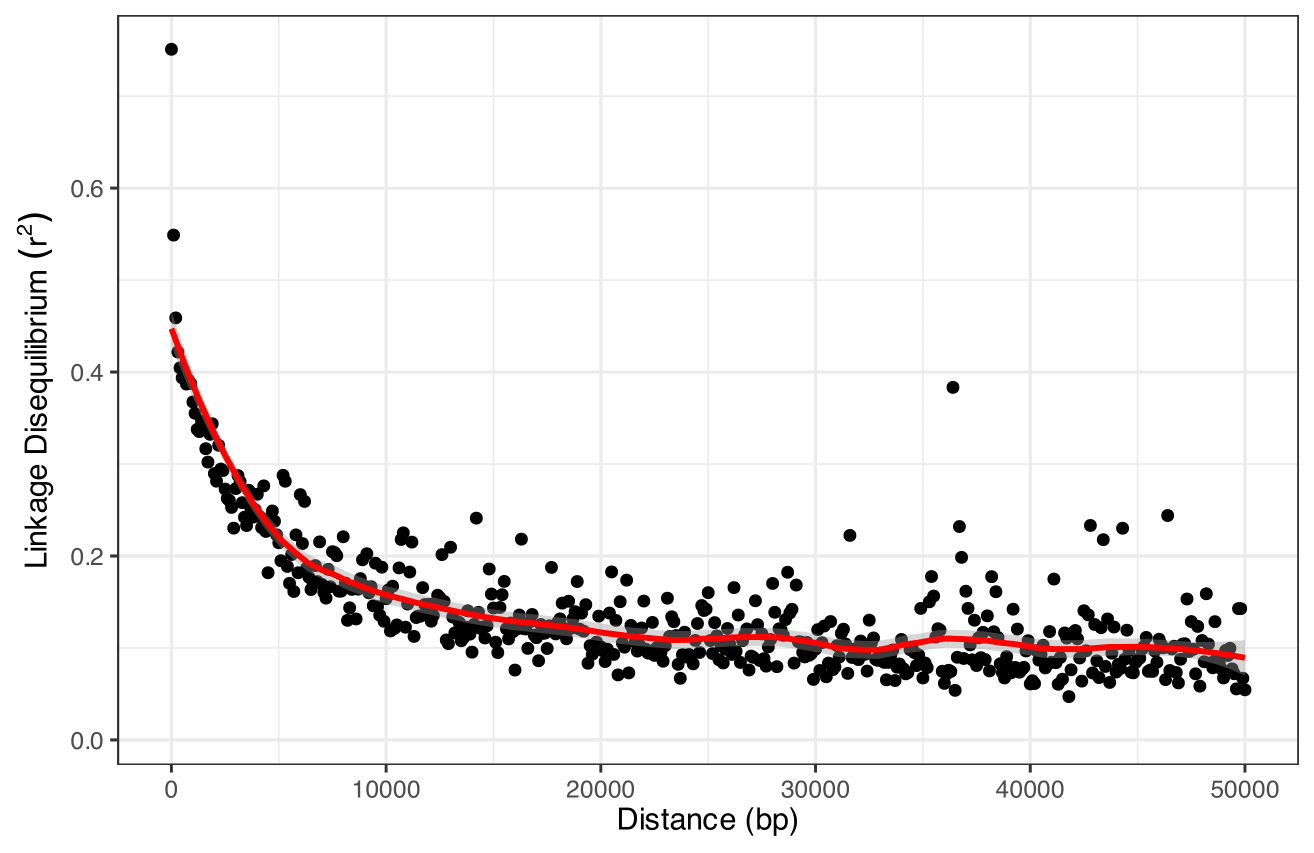

**Figure S13. Linkage disequilibrium (LD) decay across physical distance.** LD was measured as the squared correlation coefficient (r²) between SNP pairs with physical distances less than 100 kb. SNP pair distances were binned into 100 bp intervals, and mean r² values were calculated within each bin to summarize LD decay. A locally estimated scatterplot smoothing (LOESS) curve (red line, span = 0.3) was fitted to the binned mean r² values (black points) to visualize the trend of LD decay with increasing distance.

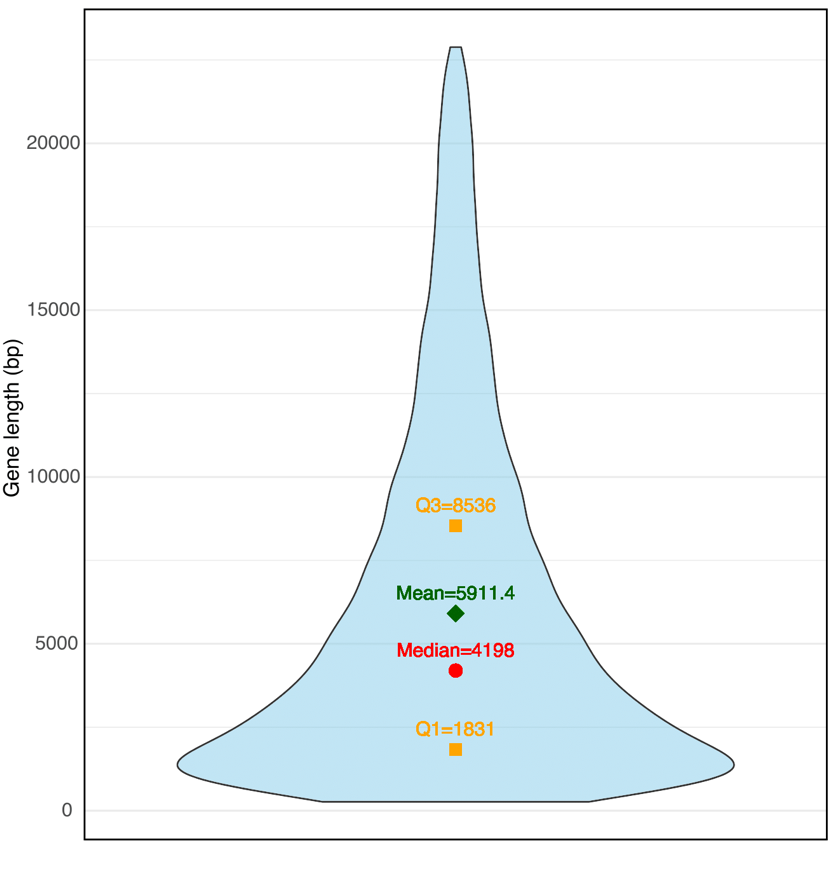

**Figure S14.** **Gene lengths (in base pairs) density distribution in the annotated *S. pistillata* reference genome.** The largest 5% of genes were removed to reduce the influence of extreme outliers. Key summary statistics highlight the median, mean, first and third quartiles of the distribution. The majority of genes fall between ~1.8 kb and ~8.5 kb.

**Table S1. Detailed information on sample sizes per location.** For each population (identified by genomic PCA), we report the reef name, reef site (which include habitat information: Back reef slope, Front reef slope, Flank reef slope and lagoon), depth zone (shallow or deep), number of samples retained after quality filtering, and geographic coordinates.

| Population | Reef | Site | Depth | N samples | Latitude | Longitude |
| --- | --- | --- | --- | --- | --- | --- |
| Heron | Heron | Back1 | Deep | 1 | -23.428 | 151.952 |
| Heron | Heron | Back1 | Shallow | 2 | -23.428 | 151.952 |
| Heron | Heron | Back2 | Deep | 4 | -23.434 | 151.92 |
| Heron | Heron | Back2 | Shallow | 3 | -23.434 | 151.921 |
| Heron | Heron | Flank1 | Shallow | 5 | -23.456 | 151.926 |
| Heron | Heron | Front1 | Shallow | 1 | -23.472 | 151.978 |
| Heron | Heron | Front2 | Deep | 5 | -23.471 | 151.951 |
| Heron | Heron | Front2 | Shallow | 1 | -23.47 | 151.951 |
| Lady Musgrave | Lady Musgrave | Flank2 | Shallow | 7 | -23.918 | 152.392 |
| Lady Musgrave | Lady Musgrave | Lagoon1 | Shallow | 1 | -23.896 | 152.414 |
| Lizard | Lizard | Back1 | Shallow | 3 | -14.651 | 145.45 |
| Lizard | Lizard | Back2 | Shallow | 6 | -14.668 | 145.442 |
| Lizard | Lizard | Flank1 | Shallow | 3 | -14.698 | 145.445 |
| Lizard | Lizard | Front1 | Shallow | 8 | -14.691 | 145.469 |
| Lizard | Lizard | Front2 | Shallow | 7 | -14.649 | 145.493 |
| Lizard | Lizard | Lagoon1 | Shallow | 5 | -14.688 | 145.465 |
| Moore | Moore | Back1 | Deep | 5 | -16.847 | 146.217 |
| Moore | Moore | Back2 | Deep | 4 | -16.882 | 146.184 |
| Moore | Moore | Back2 | Shallow | 1 | -16.882 | 146.184 |
| Moore | Moore | Front1 | Deep | 4 | -16.872 | 146.254 |
| Moore | Moore | Front1 | Shallow | 1 | -16.872 | 146.254 |
| Moore | Moore | Front2 | Deep | 5 | -16.883 | 146.245 |
| Moore | Moore | Front2 | Shallow | 4 | -16.883 | 146.245 |
| Moore | Moore | Lagoon1 | Deep | 4 | -16.86 | 146.226 |
| Moore | Moore | Lagoon1 | Shallow | 5 | -16.86 | 146.226 |
| Central | Chicken | Back 1 | Shallow | 9 | -18.653 | 147.709 |
| Central | Chicken | Back 2 | Shallow | 4 | -18.66 | 147.703 |
| Central | Davies | Back 1 | Deep | 3 | -18.826 | 147.627 |
| Central | Davies | Back 1 | Shallow | 9 | -18.826 | 147.627 |
| Central | Davies | Back 2 | Shallow | 9 | -18.814 | 147.637 |
| Central | Davies | Flank 1 | Shallow | 11 | -18.806 | 147.669 |
| Central | Davies | Flank1 | Deep | 8 | -18.806 | 147.669 |
| Central | Davies | Front 1 | Deep | 7 | -18.82 | 147.664 |
| Central | Davies | Front 1 | Shallow | 4 | -18.819 | 147.664 |
| Central | Davies | Front 2 | Shallow | 7 | -18.832 | 147.653 |
| Central | Davies | Front 2 | Deep | 7 | -18.832 | 147.653 |
| Central | Davies | Lagoon 1 | Shallow | 12 | -18.833 | 147.633 |
| Central | Davies | Lagoon 2 | Shallow | 4 | -18.831 | 147.633 |
| Central | Little Broadhurst | Back 1 | Shallow | 8 | -18.949 | 147.693 |
| Central | Little Broadhurst | Flank 1 | Shallow | 6 | -18.944 | 147.699 |
| Central | Little Broadhurst | Lagoon 1 | Shallow | 6 | -18.949 | 147.702 |
| Pelorus | Pelorus | Back 1 | Deep | 3 | -18.551 | 146.488 |
| Pelorus | Pelorus | Back 2 | Deep | 1 | -18.541 | 146.488 |
| Pelorus | Pelorus | Back 2 | Shallow | 3 | -18.541 | 146.489 |
| Pelorus | Pelorus | Flank 1 | Deep | 4 | -18.572 | 146.495 |
| Pelorus | Pelorus | Front 1 | Deep | 10 | -18.539 | 146.5 |

**Table S2. Functional features associated with the eleven windows repeatedly associated with temperature variation across populations (PicMin *q*<0.1).** For each 10-kb genomic window, we report the associated scaffold, gene coordinates, and details of any overlapping annotated genes. Gene biotype and functional annotation are derived from the NCBI Eukaryotic Genome Annotation Pipeline (egapx v0.5.2) and uncharacterized protein-coding genes were subject to additional AlphaFold and FoldSeek searches.

| Scaffold | Gene ID | Biotype | Annotation |  |
| --- | --- | --- | --- | --- |
| JAUZEC010000071.1 | egapxtmp_000003 | protein-coding | Leukocyte elastase inhibitor-like | — |
| JAUZEC010000071.1 | egapxtmp_030525 | lncRNA | — | — |
| JAUZEC010000071.1 | egapxtmp_000004 | protein-coding | Stimulated by retinoic acid gene 6 protein-like | — |
| JAUZEC010000102.1 | egapxtmp_028464 | lncRNA | — | — |
| JAUZEC010000380.1 | egapxtmp_024018 | protein-coding | Uncharacterized | Transcobalamin-like C-terminal domain-containing protein |
| JAUZEC010000380.1 | egapxtmp_024017 | protein-coding | Uncharacterized | Transcobalamin-like C-terminal domain-containing protein |
| JAUZEC010001083.1 | egapxtmp_005275 | protein-coding | Peroxidasin homolog | — |
| JAUZEC010001110.1 | egapxtmp_012096 | protein-coding | NF-X1-type zinc finger protein NFXL1-like | — |
| JAUZEC010001110.1 | egapxtmp_010025 | protein-coding | Uncharacterized | Peptidase C1A papain C-terminal domain-containing protein |
| JAUZEC010001113.1 | egapxtmp_012844 | protein-coding | Sulfotransferase family protein | — |
| JAUZEC010001548.1 | egapxtmp_004370 | protein-coding | Uncharacterized | Sulfotransferase family protein |
| JAUZEC010001548.1 | egapxtmp_004577 | protein-coding | Uncharacterized | Sulfotransferase family protein |
| JAUZEC010001548.1 | egapxtmp_004371 | protein-coding | Uncharacterized | Transcobalamin-like C-terminal domain-containing protein |
| JAUZEC010003464.1 | egapxtmp_020407 | protein-coding | Uncharacterized | **Ig-like domain-containing protein** |
| JAUZEC010003578.1 | egapxtmp_022060 | protein-coding | Basement membrane-specific heparan sulphate proteoglycan core protein-like |  |

**Table S3. Proportion of genetic variation explained by environmental predictors in RDA models across datasets.** For each model, we report the degrees of freedom for the constrained axes (i.e., Df, the number of environmental predictors in your RDA model), the model significance (*p*-value from permutation tests), the proportion of total genomic variance explained by all predictors in the RDA model (raw: R², and adjusted for number of predictors and sample size: adj R²), followed by the percentage of relative contributions for each environmental or geographic predictor to the explained genomic variation (summing to 1 for each model).

| **Population** | **Df** | ***p*-value** | **R²** | **adj R²** | **env-PC1** | **env-PC2** | **env-PC3** |  | **Lat** | **Long** |
| --- | --- | --- | --- | --- | --- | --- | --- | --- | --- | --- |
| **GBR-wide Environment** | 3 | 0.001 | 0.040 | 0.030 | 0.520 | 0.330 | 0.150 |  | — | — |
| **GBR-wide Geography** | 2 | 0.001 | 0.037 | 0.028 | — | — | — |  | 0.62 | 0.38 |
| **GBR-wide Partial** | 3 | 0.001 | 0.026 | 0.014 | 0.60 | 0.21 | 0.19 |  | — | — |
| **Lizard** | 3 | 0.001 | 0.110 | 0.012 | 0.360 | 0.330 | 0.310 |  | — | — |
| **Moore** | 3 | 0.001 | 0.100 | 0.011 | 0.380 | 0.320 | 0.300 |  | — | — |
| **Central** | 3 | 0.001 | 0.030 | 0.060 | 0.380 | 0.340 | 0.280 |  | — | — |
| **Pelorus** | 3 | 0.001 | 0.160 | 0.014 | 0.350 | 0.340 | 0.310 |  | — | — |
| **Heron** | 3 | 0.001 | 0.150 | 0.009 | 0.370 | 0.330 | 0.310 |  | — | — |

**Table S4.** **Pearson correlation coefficients among the six environmental variables used in redundancy analyses.** Pairwise Pearson’s *r* values between mean photosynthetically active radiation (PAR), mean turbidity, mean bottom water velocity, distance to shore, mean temperature and monthly temperature range, across the sampling range (i.e., GBR-wide).

|  | **PAR** | **Mean turbidity** | **Mean velocity** | **Distance to shore** | **Mean temp** | **Temp range** |
| --- | --- | --- | --- | --- | --- | --- |
| **PAR** | 1.00 |  |  |  |  |  |
| **Mean turbidity** | -0.35 | 1.00 |  |  |  |  |
| **Mean velocity** | 7.3e-02 | -0.12 | 1.00 |  |  |  |
| **Distance to shore** | 0.40 | -0.52 | -0.34 | 1.00 |  |  |
| **Mean temp** | 0.16 | -0.42 | 8.3e-03 | -0.34 | 1.00 |  |
| **Temp range** | -4.8e-02 | -0.19 | 1.00 | -0.23 | -0.19 | 1.00 |
